# N^6^-methyladenosine mRNA marking promotes selective translation of regulons required for human erythropoiesis

**DOI:** 10.1101/457648

**Authors:** Daniel A. Kuppers, Sonali Arora, Yiting Lim, Andrea Lim, Lucas Carter, Philip Corrin, Christopher L. Plaisier, Ryan Basom, Jeffrey J. Delrow, Shiyan Wang, Housheng Hansen He, Beverly Torok-Storb, Andrew C. Hsieh, Patrick J. Paddison

## Abstract

Many of the regulatory features governing erythrocyte specification, maturation, and associated disorders remain enigmatic. To identify new regulators of erythropoiesis, we performed a functional genomic screen for genes affecting expression of the erythroid marker CD235a/GYPA. Among validating hits were genes coding for the N^6^-methyladenosine (m^6^A) mRNA methyltransferase (MTase) complex, including, *METTL14*, *METTL3*, and *WTAP*. We found that m^6^A MTase activity promotes erythroid gene expression programs and lineage specification through selective translation of >200 m^6^A marked mRNAs, including those coding for SETD methyltransferase, ribosome, and polyA RNA binding proteins. Remarkably, loss of m^6^A marks resulted in dramatic loss of H3K4me3 across key erythroid-specific KLF1 transcriptional targets (e.g., Heme biosynthesis genes). Further, each m^6^A MTase subunit and a subset of their mRNAs targets, including *BRD7*, *CXXC1*, *PABPC1*, *PABPC4*, *STK40*, and *TADA2B*, were required for erythroid specification. Thus, m^6^A mRNA marks promote the translation of a network of genes required for human erythropoiesis.

## Main

In adult humans, erythropoiesis occurs in a step-wise lineage progression from bi-potent MEP to proliferative erythroid progenitors to mature erythrocytes to fulfill daily requirement for ~2×10^11^ new erythrocytes^1^. The complex process of erythroid lineage commitment and maturation is governed by multiple regulatory mechanisms, including: a) transcriptional, epigenetic, and RNAi-dependent promotion of lineage-specific cell characteristics and restriction of developmental potential^2,3^; b) lineage-specific cytokine signaling, most notably erythropoietin (EPO) and its receptor, that promote cell survival and expansion of committed progenitors^4^; and c) a network of ribosome-associated proteins (e.g., *RPS19*), which when mutated can trigger life-threatening anemias (e.g., Diamond-Blackfan anemia (DBA)) and myeloproliferative disease by blocking erythroid maturation^5^. However, owing to the limitations of deriving and manipulating human hematopoietic stem and progenitor cells (HSPCs) and erythroid progenitors, experimental investigations of erythropoiesis have been limited in scope. As a result, many additional regulatory factors governing human erythropoiesis likely await discovery.

N6-methyladenosine (m^6^A) is an abundant modification of mRNA with an increasingly important role in normal cell physiology^6–10^ and disease^11–15^. The core m^6^A methyltransferase complex (MTase) consists of a trimeric complex containing two proteins with conserved MTase (MT-A70) domains, METTL3 and METTL14^16,17^, and an additional subunit, the Wilms’ tumor 1-associating protein (WTAP)^18,19^. In cell-based models, m^6^A has been shown to participate in numerous types of mRNA regulation, including pre-mRNA processing^17,20,21^, mRNA translation efficiency^22^, mRNA stability^16^, and miRNA biogenesis^23^. In the context of hematopoiesis, m^6^A has recently been found to regulate the expansion and self-renewal of hematopoietic stem cells^24,25^, as well as functioning as a negative regulator of myelopoiesis and a potential driver of acute myeloid leukemia (AML)^11,26^.

In this study, we utilized a comprehensive CRISRP-Cas9-based screening approach to identify an essential regulatory role for m^6^A RNA methylation during erythropoiesis ^27^. Loss of m^6^A through inhibition of the methyltransferase complex results in disruption of the erythroid transcriptional program without direct inhibition of previously identified master transcriptional regulators (e.g. GATA1, KLF1). Instead, we observe translational down regulation of a variety of genes with known or suspected roles in erythropoiesis, erythroid-related diseases, and/or hematopoietic progenitor cell function. In addition, the transcriptional changes are in part driven by m^6^A translational regulation of the SETD1A/B complex, resulting in loss of the transcriptional activation mark, H3K4me3, when the m^6^A MTase complex is inhibited. Furthermore, these findings have potentially important implications for understanding myelodysplastic syndromes (MDS) as well as certain anemias, and they present a novel mechanism for the regulation of histone epigenetics.

## Results

### Validation of HEL cells as a surrogate model of erythropoiesis for whole genome CRISPR screening

Since technical limitations precluded the use of human HSPCs for large scale functional genomic screens, we took advantage of HEL cells as a surrogate model, which express key markers and transcription factors associated with erythropoiesis^28^. We chose CD235a as the screen readout, since this cell surface marker is the major sialoglcyoprotein found on the erythrocyte membrane and is a faithful indicator of erythroid lineage maturation^29^. We first performed pilot studies with sgRNAs targeting *GYPA*, which encodes CD235a, and two key transcriptional regulators of *GYPA* expression, GATA1 and LMO2^2,3,30^. Transduction of HEL cells with lentiviral (lv) vectors expressing sgRNAs targeting *GYPA*, *GATA1*, or *LMO2* resulted in significant reduction of CD235a expression (Supplementary Fig. 1a) and, for *sgGATA1* and *sgLMO2*, significant changes in gene expression of key erythroid gene targets, including, *ALAS2*, *EPOR*, *GYPA*, and *KLF1*, as well as erythroid stage-specific gene expression programs (Supplementary Fig. 1b,c). The results indicated that HEL cells can recapitulate at least a portion of the molecular features associated with erythropoiesis.

### CRISPR-Cas9 whole genome screening in HEL cells to identify regulators of erythropoiesis

Next, we performed a pooled lv-based CRISPR-Cas9 screen targeting 19,050 genes and 1,864 miRNAs^27^ in HEL cells using two rounds of antibody-based CD235a+ cell depletion to derive a population of CD235a-/low cells (day 12 post-infection) (Fig. 1a, and Supplementary Fig. 1d). CD235a-/low cells were then subjected to sgRNA-seq^31^ to identify sgRNAs enriched in this population compared to the starting population and, also, cells which were outgrown in culture for 12 days (Fig. 1a, and Supplementary Fig. 1e, Supplementary Table 1). Using filter criteria shown in Supplementary Fig. 1f, we identified 31 candidate genes which were then individually retested with lv-sgRNAs for modulation of CD235a expression. A total of 12 genes retested in HEL cells (Fig. 1b), primarily falling into 3 categories: components of the glycosylation machinery^32^; a modification required for antibody recognition of CD235a^33^, key components of the GATA1 transcriptional regulatory complex^2,3^; and, surprisingly, the trimeric m^6^A mRNA MTase^34,35^ complex, *METTL14*, *METTL3*, and *WTAP* (Fig. 1b-d). Given this potential new role for m^6^A-dependent regulation of erythropoiesis, we further examined the underlying biology and phenotypes associated with these latter hits.

**Fig. 1:**
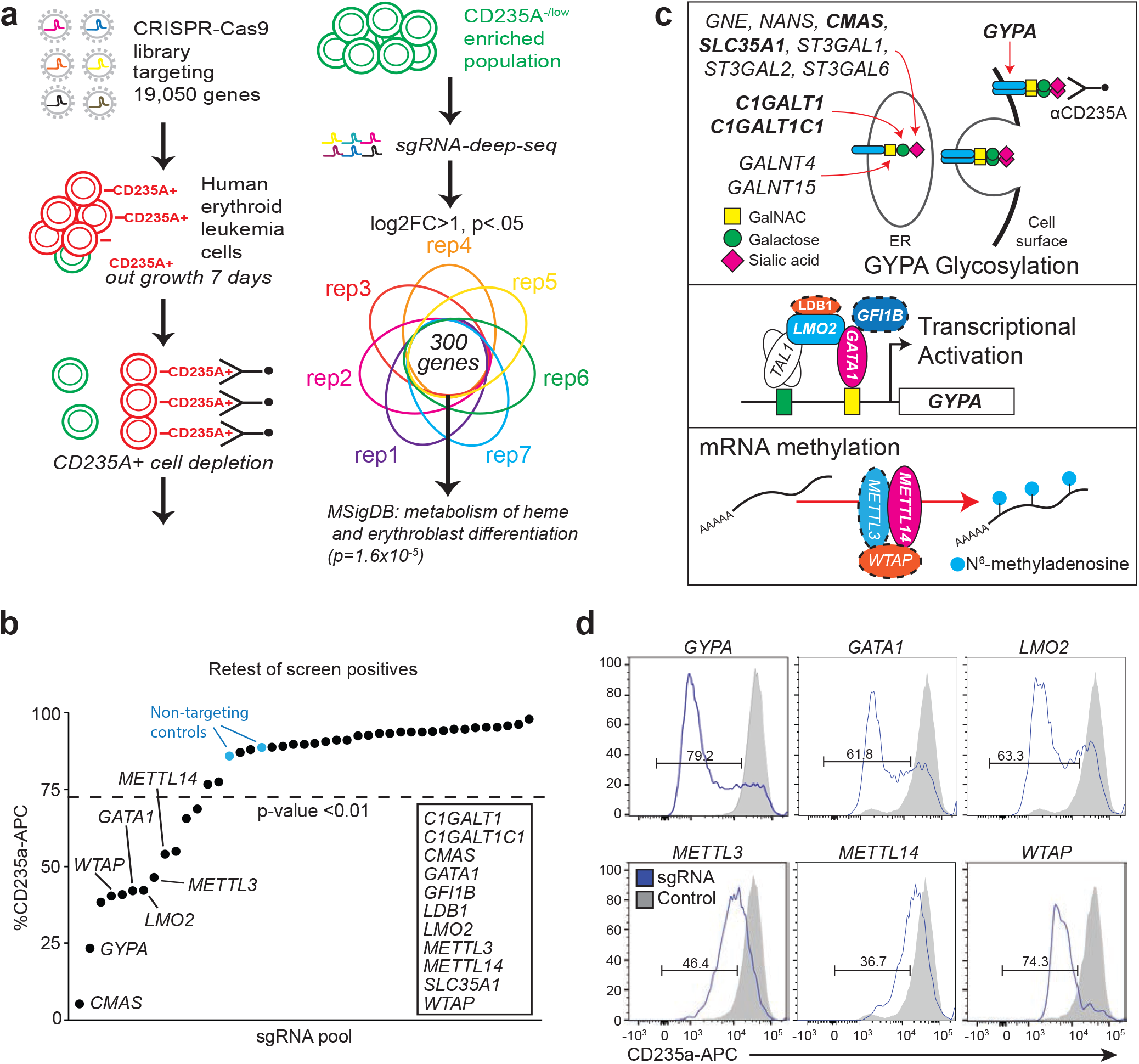
CRISPR-Cas9 whole genome screening in HEL cells to identify regulators of erythropoiesis. **a,** CRISPR-Cas9 screen design for enrichment of sgRNAs promoting the CD235a-/low state. **b,** Individual genes retested by flow cytometry for CD235a surface expression on day 10 posttransduction in HEL cells. Cells were transduced with individual gene retests pools of 4 sgRNAs (see Methods for details). A modified z-score cutoff for a p-value<0.01 was used to define a positive hit, with all scoring genes indicated within the box. For the m^6^A MTase complex, only METTL14 scored in the primary screen. We found that the sgRNAs targeting METTL3 and WTAP in the screening library were not effective and substituted sgRNAs from the human CRISPR Brunello lentiviral pooled library for METTL3 and WTAP. LDB1 and GFI1B each scored with 1 sgRNA in the initial screen and new sgRNAs were generated, also from the Brunello library. **c,** Diagram of the three primary categories of screen hits. Top panel shows genes with multiple sgRNA hits in bold; all others have a single sgRNA scoring. Middle and bottom panels: genes validated by secondary individual gene tests highlighted in color; solid lines indicate 2 or more sgRNAs scored from primary screen, while dashed lines indicate 0 or 1 sgRNAs scored from primary screen. **d,** Representative flow cytometry results for positive retest hits in HEL cells. Cells were transduced with lentiCRISPRv2-mCherry virus and assayed by FACS 7-9 days later.

### m^6^A marks the mRNA of key hematopoietic and erythroid regulators

To first reveal how m^6^A mRNA marks affect erythroid regulatory networks, we performed methylated RNA immunoprecipitation sequencing (MeRIP-seq)^36,37^, which provides a quantitative site-specific readout of all m^6^A-modified transcripts, and in parallel, examined changes in gene expression, and mRNA splicing (Fig. 2,3 Supplementary Fig. 2,3). Profiling the poly-A RNA m^6^A methylome of HEL cells revealed a total of 19,047 m^6^A peaks in 7,266 protein coding genes, representing 42.7% of genes expressed in HEL cells (Supplementary Table 2). The number of m^6^A peaks per gene ranged as high as 28, with 64.3% of m^6^A containing mRNAs having one or two peaks (Supplementary Fig. 2a). Consistent with previous MeRIP-seq results^36,37^, we observed enrichment of peaks around the stop codon of protein-coding mRNAs and a similar adenosine methylation site motif of GAACU, compared to the previously identified “RRACH”^37^ (Fig. 2a,b). Critically, m^6^A-marked mRNAs in HEL cells were enriched for key regulators of hematopoiesis and erythropoiesis (e.g., *GATA1*, *FLI1*, *KLF1*, and *MPL*) (Fig. 2c) and genes with causal roles in erythroid-related diseases (Fig. 2d,e). The same m^6^A-marked mRNAs were not observed in human embryonic kidney cells (Supplementary Fig. 2b,c).

**Fig. 2:**
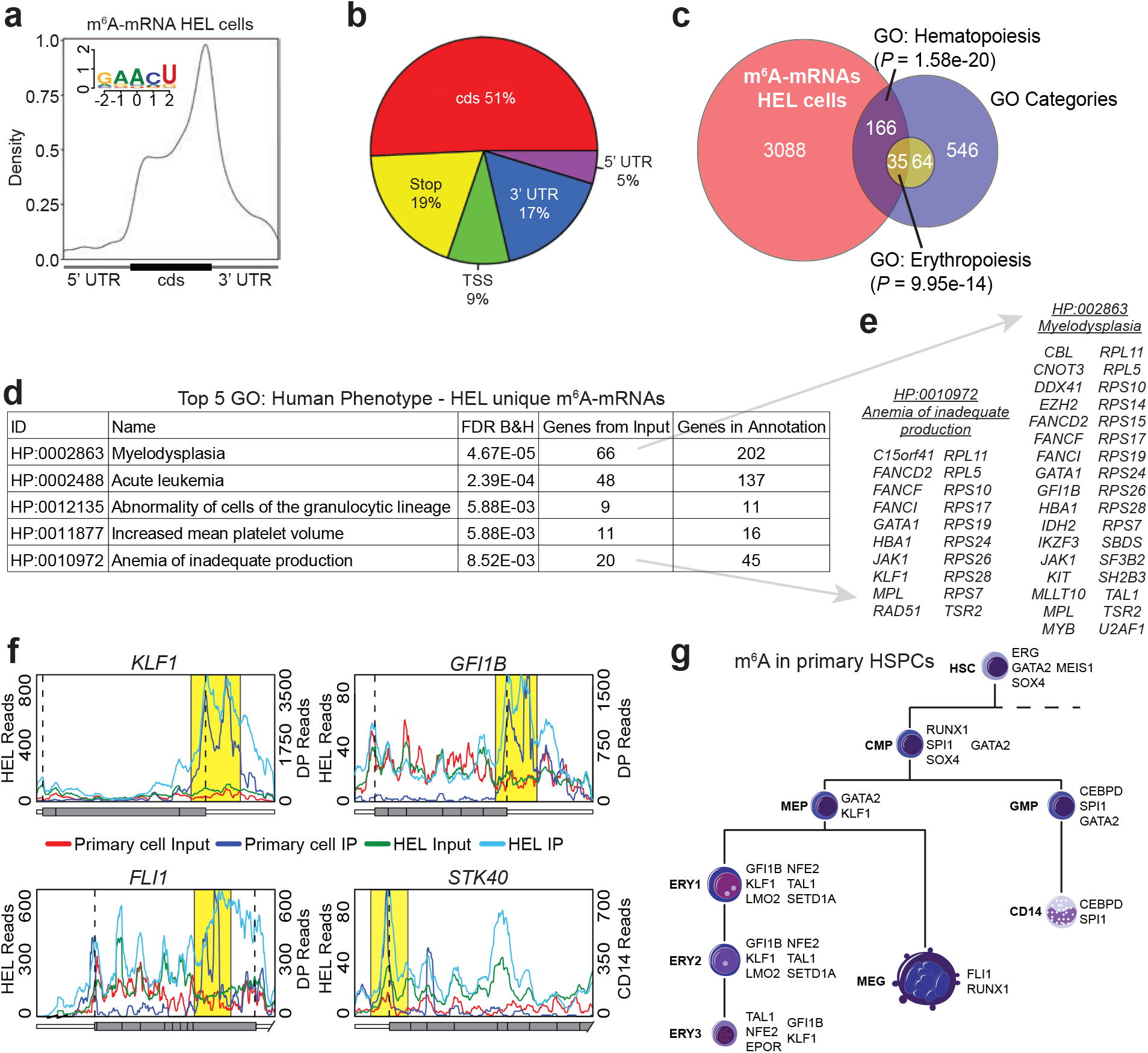
m^6^A marks the mRNA of key hematopoietic and erythroid regulators. **a,** Distribution of m^6^A sites detected by meRIP-seq in HEL cells (150 μg), with an enrichment around the stop site, enriched m^6^A methylation site motif. **b,** Pie chart displaying the frequency of m^6^A peaks, from HEL (150 μg), within different transcript regions: TSS, centered around translation start ATG, Stop, centered around the stop codon. **c,** Genes uniquely methylated in HEL cells versus 293T cells are enriched for key hematopoiesis and erythropoiesis genes. **d,** Top GO terms for HEL unique m^6^A mRNAs are enriched for genes with causal roles in hematopoietic diseases. **e,** Gene callouts for GO terms identified in panel d. **f,** Transcript maps for erythropoiesis regulators detected as methylated in both HEL (150 μg) and adult BM cells show overlapping methylation patterns (Yellow highlight). **g,** Highlighting key hematopoietic regulators detected as methylated by meRIP-seq in flow sorted adult human BM cells. The parameters used to define the hematopoietic populations are outlined in Supplementary Fig. 2i.

**Fig. 3:**
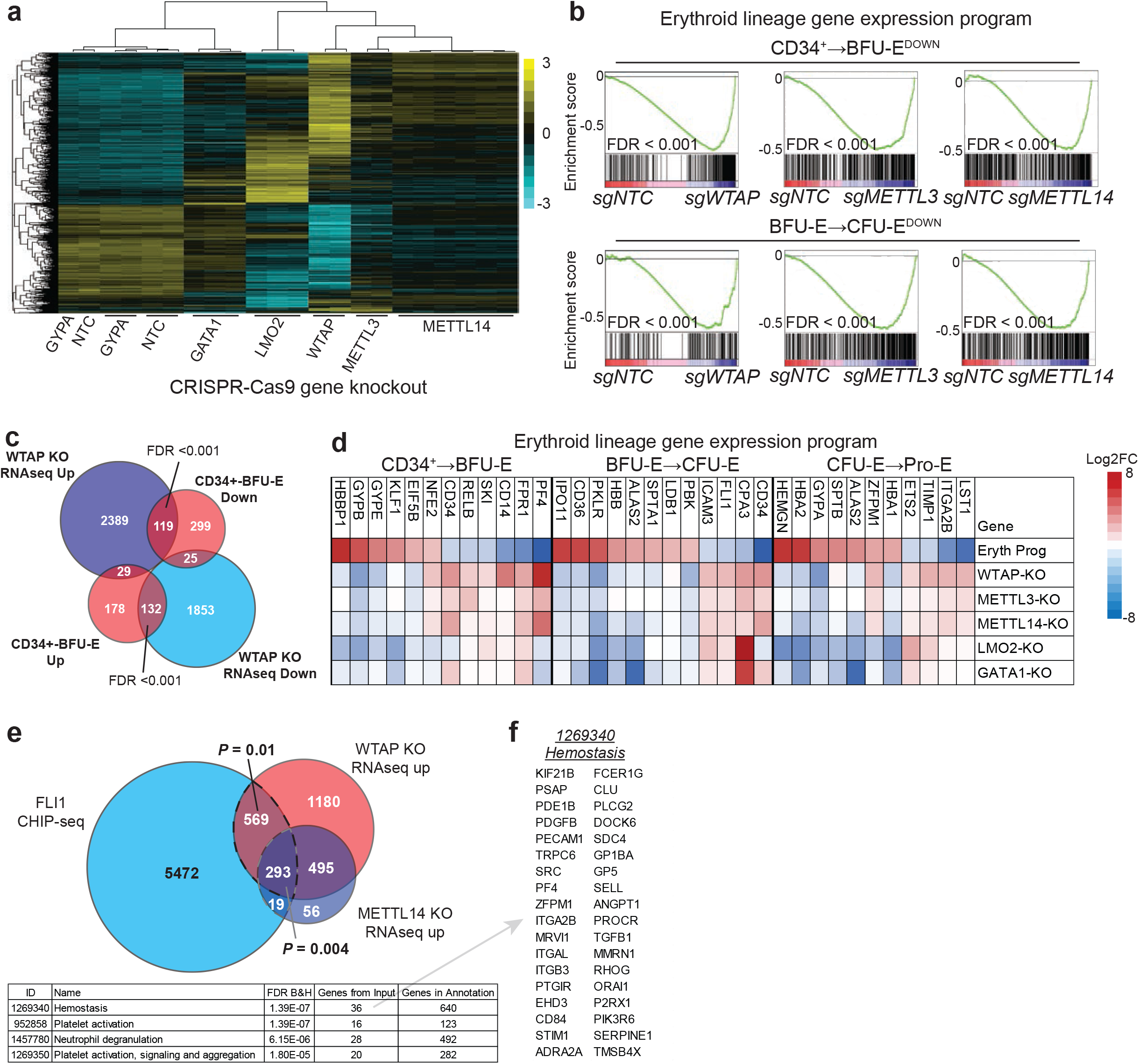
m^6^A-dependent regulation of erythroid gene expression programs. **a,** A heatmap for RNAseq analysis of HEL cells following KO of *WTAP*, *METTL3* and *METTL14*, as well as select erythroid genes, shows clustering of the three components of the m^6^A MTase complex and a unique transcriptional profile versus the GATA1 transcriptional program. **b,** GSEA analysis of *WTAP*-KO, *METTL3*-KO and *METTL14*-KO HEL cell RNA-seq, using custom gene sets for transcriptionally up and down genes during erythropoiesis as defined by^38^, shows a negative correlation between genes down regulated during erythropoiesis and transcriptional changes following m^6^A loss in HEL cells. The analysis of the additional categories can be found in Supplementary Fig. 3b. **c,** A Venn diagram of transcriptional changes in *WTAP*-KO HEL cells and transcriptional changes during early erythropoiesis shows a negative correlation. **d,** A heatmap of transcriptional changes observed during normal erythropoiesis compared to m^6^A MTase KO HEL cells and KO HEL cells for select erythroid transcriptional regulators shows similar inverse patterns of gene expression.

To ensure relevance to normal cells, we also analyzed m^6^A-marked mRNAs from 9 flow sorted hematopoietic stem and progenitor cell populations (HSPCs) from freshly harvested adult human bone marrow, including those relevant to erythropoiesis. However, given the limited quantities of mRNA which can be isolated from these populations, we took advantage of a new protocol for developed by He and colleagues for performing MeRIP-seq from as little as 2 μg of total RNA (in press, PLOS Biology), as opposed to large of total RNA quantities used for MeRIP-seq for our HEL cells (e.g., 100-300 μg)^36,37^. To validate the He protocol, we initially characterized the quality of the data by comparing the HEL cell results with the established MeRIP-seq protocol. Profiling the ribosomal RNA-depleted m^6^A methylome from 3 μg of HEL cell total RNA, we identified a total of 7,282 m^6^A peaks in 3,210 protein coding genes (Supplementary Table 2). The distribution of the number of m^6^A peaks per gene was similar between the protocols with 67.4% vs. 72.7% of m^6^A containing mRNAs having one or two peaks (Supplementary Fig. 2a,f). We also observed a similar distribution of peaks within transcripts of protein-coding mRNAs and a similar adenosine methylation site motif of GGACU (Fig. 2a,b vs Supplementary Fig. 2e,g). Overall, we observed a high level of agreement between the protocols, with a 94% overlap in methylated genes and a 76% overlap in called peaks identified with the He protocol (Supplementary Fig. 2h). However, we observe that the He protocol under samples the m^6^A methylome (Supplementary Fig. 2h).

Applying this technique to 9 HSPC populations (HSC, CMP, GMP, MEP, CD14, MEG, ERY1-3, Supplementary Fig. 2i lists the criteria used to define each population), we observe that 80.0% of the detected m^6^A methylated mRNAs in the HSPC populations overlapped with the HEL cells, with individual populations ranging from 94.8% overlap in the ERY1 population to 65.3% in the ERY3 population (Supplementary Fig. 2i). The specific m^6^A peaks detected in the HSPCs also had a high degree of overlap with those observed in the HEL cells (Fig. 2f). Consistent with the HEL cells, the m^6^A-marked mRNAs in the HSPCs were enriched for key regulators of hematopoiesis and erythropoiesis (e.g., *GATA2*, *FLI1*, *KLF1*, and *SPI1*), reinforcing the notion that m^6^A likely plays a role in regulating a variety of stages of hematopoiesis, including erythropoiesis (Fig. 2g).

### m^6^A-dependent regulation of erythroid gene expression programs

We next examined *WTAP, METTL3 and METTL14* knockout (KO) changes to steady-state mRNAs levels in HEL cells. Overall, KO of the three components of the MTase complex resulted in similar transcriptional changes, with a stronger effect observed with *WTAP* KO, a finding consistent with the more penetrant CD235a phenotype (Fig. 1d,3a). We found 4073 mRNAs significantly changed after *WTAP* KO via RNA-seq (p<.01), with 2,254 up-and 1,819 down-regulated, 1102 mRNAs significantly changed after *METTL3* KO via RNA-seq (p<.01), with 687 up and 415 down-regulated, and 1505 mRNAs significantly changed after *METTL14* KO via RNA-seq (p<.01), with 993 up and 512 down-regulated.

Examining overlap with m^6^A marked mRNAs revealed for up regulated mRNAs the m^6^A frequency did not deviate from HEL cells (39.6% overall versus 38-42.8% for *METTL14*, *METTL3*, or *WTAP* KO). However, for *WTAP* and *METTL3* KOs we observed significant enrichment for m^6^A marking among down regulated mRNAs (*WTAP* 55.0%, pval=0.008; *METTL3*, 52.8%, pval=0.009). We wondered whether this group of genes would contain key erythroid transcription factors that promote *GYPA* expression, as levels of *GYPA* mRNA, which is not m^6^A methylated, drops significantly in *WTAP, METTL3 and METTL14* KO HEL cells, similar to *GATA1* and *LMO2* KO (Fig. 3d). However, the RNA-seq data did not support this notion. Crucial *GYPA* regulators, for example, *GATA1*, *GFI1B*, *KLF1*, *LDB1*, *LMO2*, *NFE2*, and *ZFPM1/2*, were not down regulated (Supplementary Fig. 3a). Despite this, *WTAP, METTL3 and METTL14* KO significantly down regulated mRNAs associated with key stages of erythropoiesis, including, the BFU-E, CFU-E and Pro-E stages^38^ (Fig. 3b,c, Supplementary Fig. 3b), similar to *LMO2* KO (Supplementary Fig. 1b,c). Moreover, *WTAP* KO, but not *LMO2* KO, dramatically changed total m^6^A RNA levels, demonstrating that m^6^A MTase is inhibited in *WTAP* KO cells (Supplementary Fig. 2d).

Further, we also examined changes in expression of transcriptional targets of FLI1, a pro-megakaryocyte transcription factor with inversely correlated expression to pre-erythroid transcription factor KLF1^39^. We compared promoters occupied by FLI1 in primary human megakaryocytes^40^ to our RNAseq data and found significant up regulation of these genes in HEL cells after *METTL14*, *METTL3*, and *WTAP* KO.These included key megakaryocyte and platelet-related genes, including those involved in hemostasis such as: GP1BA, PF4, and GP5 (Fig. 3e,f).

Taken together, these results suggested that m^6^A MTase activity may affect expression of erythroid genes independently of affecting steady-state mRNA levels of key erythroid transcriptional regulators and that m^6^A MTase activity suppresses expression of megakaryocytic lineage genes in HEL cells.

### m^6^A-mRNA marking does not affect splicing in *cis* in HEL cells

We next wondered whether *WTAP, METTL3 and METTL14* KO-driven changes in erythroid gene expression could be due to alteration in splicing of key erythroid transcription or other factors. It was previously asserted that m^6^A marks may promote exon inclusion in certain mRNAs^17,20^. To this end, we examined changes in all predicted exon-exon and exon-intron boundaries (using MISO^41^) and compared any altered splicing patterns to m^6^A-marked transcripts. We found that *WTAP, METTL3 and METTL14* KO resulted in 434, 226, and 306 significant splicing changes in 336, 196, and 246 genes respectively. However, we did not observe significant overlap between m^6^A peaks at exon/intron boundaries where *WTAP, METTL3 and METTL14* KO-induced splicing alterations occurred, either in general or for specific classes of splicing events (Supplementary Fig. 3c) and the changes were mostly modest (59.5% median delta psi< ±0.25)( Supplementary Table 4). This suggested that m^6^A-induced changes in splicing were not driving our phenotypes.

### m^6^A-dependent translational regulation of known or putative erythropoiesis factors

We next asked with m^6^A marking might impact the translation of key genes responsible for promoting erythroid gene expression. Since *WTAP* KO produced the most penetrant loss of CD235a among the three m^6^A MTase subunits (Fig. 1b,d), had the greatest transcriptional effect (Fig. 3a) and we demonstrated that *WTAP* is essential for m^6^A MTase activity (Supplementary Fig. 2d), all proceeding translational HEL studies were performed with *WTAP* KO. We utilized ribosome profiling, a whole transcriptome method to measure translation efficiency^42,43^, to analyze *WTAP*-dependent effects on mRNA translation. This analysis yielded 1055 genes that were translationally changed without significant alterations to mRNA levels following *WTAP* KO, with 738 up and 317 down regulated (Fig. 4a and Supplementary Table 5). Of these, ~45% of up and ~63% of down regulated mRNAs were m^6^A methylated. Interestingly, more heavily m^6^A-marked mRNAs show significantly lower mRNA ribosome association after *WTAP* KO (Supplementary Fig. 4a), indicating that m^6^A marks are directly promoting translation of a subset of mRNAs. Supporting this notion, we did not observe global changes in de novo protein synthesis in *WTAP* KO cells, as determined by puromycin incorporation of nascent peptide chains (Supplementary Fig. 4b). Intriguingly, among translationally down-regulated m^6^A-mRNAs after *WTAP* KO, we observed significant enrichment for proteins with RNA binding and histone methyltransferase activity (Supplementary Table 5), including ribosomal proteins with causal roles in human Diamond-Blackfan anemia^5,44^, and key H3K4 MTases and associated proteins, including catalytic MTases KMT2D/MLL4^45^, SETD1A^46^, and SETD1B^47^ (Fig. 4a). Furthermore, at least 48 m^6^A regulated mRNAs in the translationally down category have known or suspected roles in erythropoiesis, erythroid-related diseases, and/or hematopoietic progenitor cell function, for example, MCL1^48^, FBXW7^49,50^, and KMT2D^51,52^ (Fig. 4b and Supplementary Table 5).

**Fig. 4:**
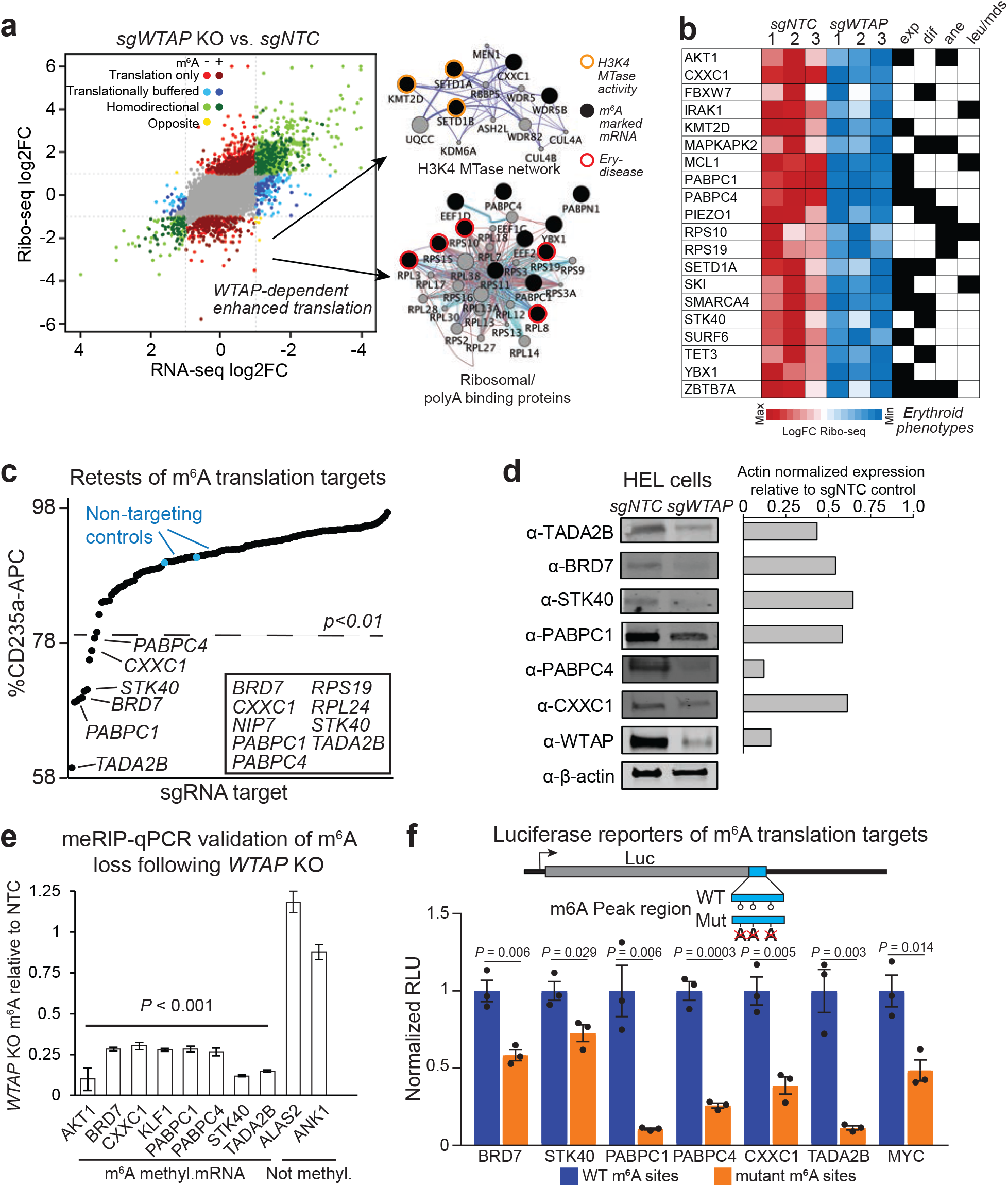
m^6^A-dependent translation regulation of known or putative erythropoiesis regulators. **a,** Translational and transcriptional changes in *WTAP*-KO HEL cells measured by ribosome profiling, network maps for interactions enriched in m^6^A uniquely translationally down genes highlight targeting of the H3K4 MTase complex, as well as ribosomal proteins. **b,** A heatmap of genes uniquely translationally down with known or suspected roles in hematopoietic progenitor cell function (exp), erythropoiesis (dif), anemia (ane), and/or other hematopoietic diseases (leu/mds). These and other genes in this category, along with associated references are in Supplementary Table 5. **c,** Several m^6^A targets, which are only translationally changed following *WTAP*-KO, can partially recapitulate the m^6^A-MTase inhibition phenotype in HEL cells when individually targeted by sgRNAs or shRNAs. Several genes that scored as “essential” in HEL cells outgrowth CRISPR-Cas9 screen (See Supplementary Table 1) were targeted with shRNAs rather than sgRNAs, including: *RPL3*, *RPL24*, *RPS19*, *RRS1*, *RUVBL2*, and *SFPQ*. Cells were infected with pools of 4 lv-sgRNA or individual shRNAs. A z-score cutoff for a p-value<0.01 was used to define a positive hit, with all scoring genes indicated within the box. **d,** Western blot validation of non-essential retest hits following *WTAP* KO in HEL cells shows reduced protein expression for all hits. **e,** meRIP-qPCR validation of altered m^6^A levels for genes translationally decreased following *WTAP* KO relative to NTC m^6^A levels. (n=3 technical replicates, mean ± SEM. t-test 2-sided) **f,** Luciferase reporter validation of translationally downregulated genes containing an m^6^A site following *WTAP* KO. The region identified by meRIP-seq as containing an m^6^A site and with the largest number of sites matching the RRACH motif were fused to the 3’ end of luciferase, except for *STK40* with a 5’ fusion, and the central A of the motif mutated. The selected region for all targets showed m^6^A dependent regulation of expression. (n=3 biological replicates, mean ± SEM. t-test 2-sided)

Since the results suggested that m^6^A-dependent translational regulation of these mRNAs could explain our erythropoiesis phenotypes, we performed functional retests on 54 genes in the translationally down category for effects on CD235a expression in HEL cells (Fig. 4c). We found that inhibition of 9 of these genes significantly decreased CD235a expression (Fig. 4c), including: *BRD7*, *CXXC1*, *NIP7*, *PABPC1*, *PABPC4*, *RPS19*, *RPL24*, *STK40*, and *TADA2B*. Critically, except for *BRD7* and *TADA2B*, each of these genes has known erythroid-related functions (Supplementary Table 5). For example, PABPC4 has been shown to bind to mRNAs associated with erythroid differentiation in mouse leukemia cells and is required for maintaining the steady-state mRNA levels of a subset of these, including CD235a/GYPA mRNA^53^, while *Stk40* deletion leads to anemia in mouse embryos characterized by a reduction in progenitors capable of erythroid differentiation^54^. Further validation of BRD7, CXXC1, PABPC1, PABPC4, STK40, and TADA2B by Western blot showed that their steady-state protein levels decreased after *WTAP* KO, supporting the ribosome profiling results (Fig. 4d). We further validated significant reduction in m^6^A levels for these genes following *WTAP* KO compared to non-m^6^A containing mRNAs by meRIP-qPCR (Fig. 4e).

To demonstrate a direct link between loss of m^6^A marking and translational down regulation, we generated luciferase reporters for BRD7, STK40, PABPC1, PABPC4, CXXC1, TADA2B and MYC, containing 197-438 bp of the m^6^A marked region with the largest number of m^6^A motifs. When tested in HEL cells, mutation of the putatively methylated A sites resulted in reduced reporter activity for all of the constructs, along with MYC which has previously been shown to be translationally regulated by m^6^A^26^ (Fig. 4f).

These results demonstrate that m^6^A marking of discrete mRNA domains promote the translation of genes with causal roles in promoting CD235a/GYPA expression and known or likely roles in erythropoiesis. The results further suggest that m^6^A mRNA regulatory elements are transferable to other mRNAs and do not depend on gene context (e.g., cis-acting chromatin binding factors specific to marked genes).

### Promoter histone H3K4me3 marks in erythroid genes are lost in the absence of m^6^A methyltransferase activity

Because we observed that WTAP activity promoted translation of a histone H3K4 MTase network, e.g., *SETD1A*, *SETD1B*, and *KMT2D* (Fig. 4a), we wondered whether H3K4 methylation would be dependent on m^6^A methyltransferase activity. In particular, previous work has shown that mice deficient for *Setd1a* show loss of promoter-associated H3K4 methylation in the erythroid lineage, loss of erythroid gene transcription, and blockade of erythroid differentiation^55^. Further, KMT2D has been shown to localize to and regulate transcription of the β-globin locus, which is specific to erythroid lineage^56^. Because the SETD1A and SETD1B have established roles in maintaining and promoting H3K4me3 and this mark is strongly associated with transcriptionally active promoters^57,58^, we focused on analysis of H3K4me3 after loss of m^6^A MTase activity. To this end, we performed CUT&RUN^59^ to map H3K4me3 marks in *METTL14, METTL3, and WTAP* KO HEL cells. By this technique, we observe 6871 H3K4me3 marked genes centered around transcription start sites (TSS) in HEL cells. Critically, we observed dramatic loss of this signal in *METTL14, METTL3, and WTAP* KO cells, with a much larger loss observed with *METTL14* and *WTAP* KOs, 1461 and 1127 H3K4me3 marked genes, respectively (Fig. 5a and c). Examples of some promoters with loss of H3K4me3 are shown in Fig. 5b, including *UROS*, a key constituent of the heme biosynthetic pathway^60^, *EPOR*, the receptor for erythropoietin, a growth factor critical for normal erythropoiesis, and *HEMGN*, a GATA1 transcriptional target^61^.

**Fig. 5:**
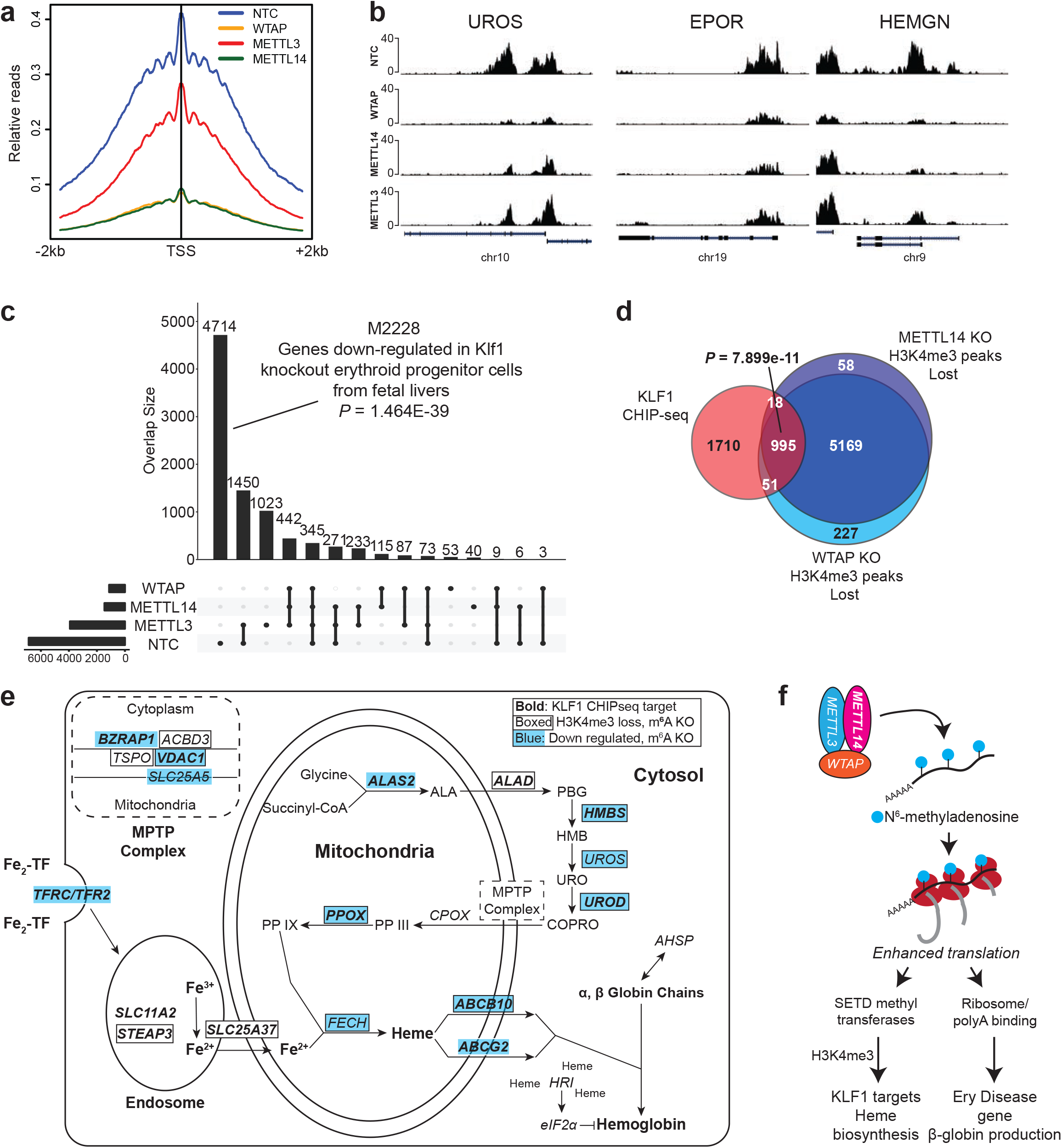
Promoter histone H3K4me3 marks are lost in the absence of m6A methyltransferase activity. **a,** An H3K4me3 relative read distribution plot around the transcription start site (TSS) of methylated genes in CRISPR NTC, *WTAP* KO, *METTL3* KO and *METTL14* KO HEL cells. The plot shows a dramatic reduction in H3K4me3 marks following loss of *WTAP* and *METTL14* with a more modest effect with loss of *METTL3*. **b,** Bedgraphs of normalized H3K4me3 CUT&RUN data for select genes with reduced methylation following m^6^A loss. **c,** An upset plot of the H3K4me3 peaks found in CRISPR NTC, *WTAP* KO, *METTL3* KO and *METTL14* KO HEL cells. The pattern of H3K4me3 loss is consistent with loss of KLF1 transcriptional regulation. Genes with reduced H3K4me3 following loss of any of the three m^6^A MTase components are enriched for genes down-regulated following *Klf1* KO. **d,** A Venn diagram showing enrichment for KLF1 CHIP-seq target genes^63^ among the H3K4me3 peaks lost following *WTAP*-KO or *METTL14*-KO, suggesting m^6^A mediated epigenetic regulation of the KLF1 transcriptional program. **e,** A diagram of the iron procurement, heme synthesis and transport, and hemoglobin assembly in erythroid cells, highlighting regulation by KLF1 and altered H3K4me3 marking following m^6^A loss. Gene highlighted in bold are KLF1 CHIPseq target genes^63^, boxed genes have reduced H3K4me3 marking following m^6^A loss, and those in blue are down regulated following m^6^A loss. These results highlight inhibition of multiple pathways involved in hemoglobin synthesis following loss of m^6^A possibly through down regulation of the KLF1 transcriptional program. **f,** The proposed model for the role of m^6^A in translational regulation of erythroid gene expression and erythropoiesis.

A more general examination of H3K4me3 loci lost after m^6^A-MTase deletion, remarkably, revealed dramatic enrichment for genes that are down regulated in mouse erythroid progenitors that have deletion of *Klf1* (GO ID M2228; P value = 1.464E-39)^62^. KLF1 is a master transcriptional regulator of erythropoiesis, facilitating many aspects of terminal erythroid differentiation including production of alpha- and beta-globin, heme biosynthesis, and coordination of proliferation and anti-apoptotic pathways^63^. However, as mentioned above, we did not observe that KLF1 was down regulated in HEL cells after *METTL14, METTL3, and WTAP* KO. So, we compared the promoters that had lost H3K4me3 after KO of m^6^A-MTase activity with those previously shown to be directly bound by KLF1 in primary erythroid progenitors^63^. Remarkably, 36% of KLF1 bound promoters lost H3K4me3 marking after inhibition of *METTL14* and *WTAP* (P value= 7.899e-11) (Fig. 5d). Further, one of the key roles of KLF1 in erythropoiesis is activating expression of genes directly involved in heme and hemoglobin biosynthesis^63,64^. We observe that m^6^A-MTase activity is required for both maintenance of promoter H3K4me4 and/or steady-state mRNA levels of most of the genes in this pathway (Fig. 5e).

In addition to KLF1 targets, we also examined H3K4me3 status of genes expressed during CD34 to BFU and CFU to ProE stages of erythropoiesis from our previous analysis (Fig. 3). We also found that a significant number of these promoters also lose H3K4me3 peaks (Supplementary Table 6), which further underscores the importance of m^6^A-mRNA regulation in promoting epigenetic marking of key erythroid genes.

Combined these data suggest a model where by m6A marks promote the translation of a broad network of key genes required for erythrocyte specification and maturation (Fig. 5f). The target genes can largely be split into two distinct pathways which when perturbed may lead to disrupted erythropoiesis: (1) H3K4me3 regulation of the KLF1 transcriptional program required for development of early erythroid progenitors and regulation of heme synthesis and hemoglobin assembly, (2) Ribosomal proteins and regulators of mRNA stability.

### Knockdown of the m^6^A MTase complex uniquely blocks erythropoiesis in human adult, bone marrow-derived HSPCs

To demonstrate that m^6^A MTase function impacts erythroid lineage specification, we knocked-down (KD) *METTL14, METTL3, and WTAP* in adult human bone marrow-derived CD34+ HSPCs with lv-shRNAs and assessed lineage formation *in vitro* (Fig. 6 and Supplementary Fig 5). In liquid cultures treated with EPO to specify erythroid lineage formation, we observed a near total loss of CD235a+ cells in m^6^A-MTase KD cells (Fig. 6a and Supplementary Fig. 5a-c). There was no observable impact on megakaryocytic differentiation (Fig. 6a and Supplementary Fig. 5a-c). For myeloid differentiation there was no observable change for *WTAP*-KD or *METTL14*-KD and an increase, though not statistically significant, from 37.5±1.8 in Scr control to 52.3±5.0 (P value=0.1062) with *METTL3*-KD (Fig. 6a and Supplementary Fig. 5a-c). Further, examination of the sequential appearance of CD71+ and CD235a+ cells during early-mid and mid-late erythroid differentiation^65^ revealed that few erythroid progenitors progressed past the earliest stages of erythropoiesis following *WTAP, METTL3 or METTL14* KD (Fig. 6b, Supplementary Fig. 5a,b). Consistent with these findings, *WTAP*-KD resulted in a complete loss of erythropoiesis in colony formation assays, including burst and colony forming units (i.e., BFU-E and CFU-E)^66^, without a noticeable effect on megakaryocyte or myeloid colony formation (Fig. 6c). However, within the MEP population (CD34+CD38+CD45RA-CD123-)^67^, *WTAP* KD did not alter the number of CD41- erythroid or CD41+ megakaryocyte committed progenitors (Supplementary Fig. 6a). Thus, the results are consistent with m^6^A-MTase activity being differentially required for early erythropoiesis in adult hematopoietic progenitors just after the MEP stage.

**Fig. 6:**
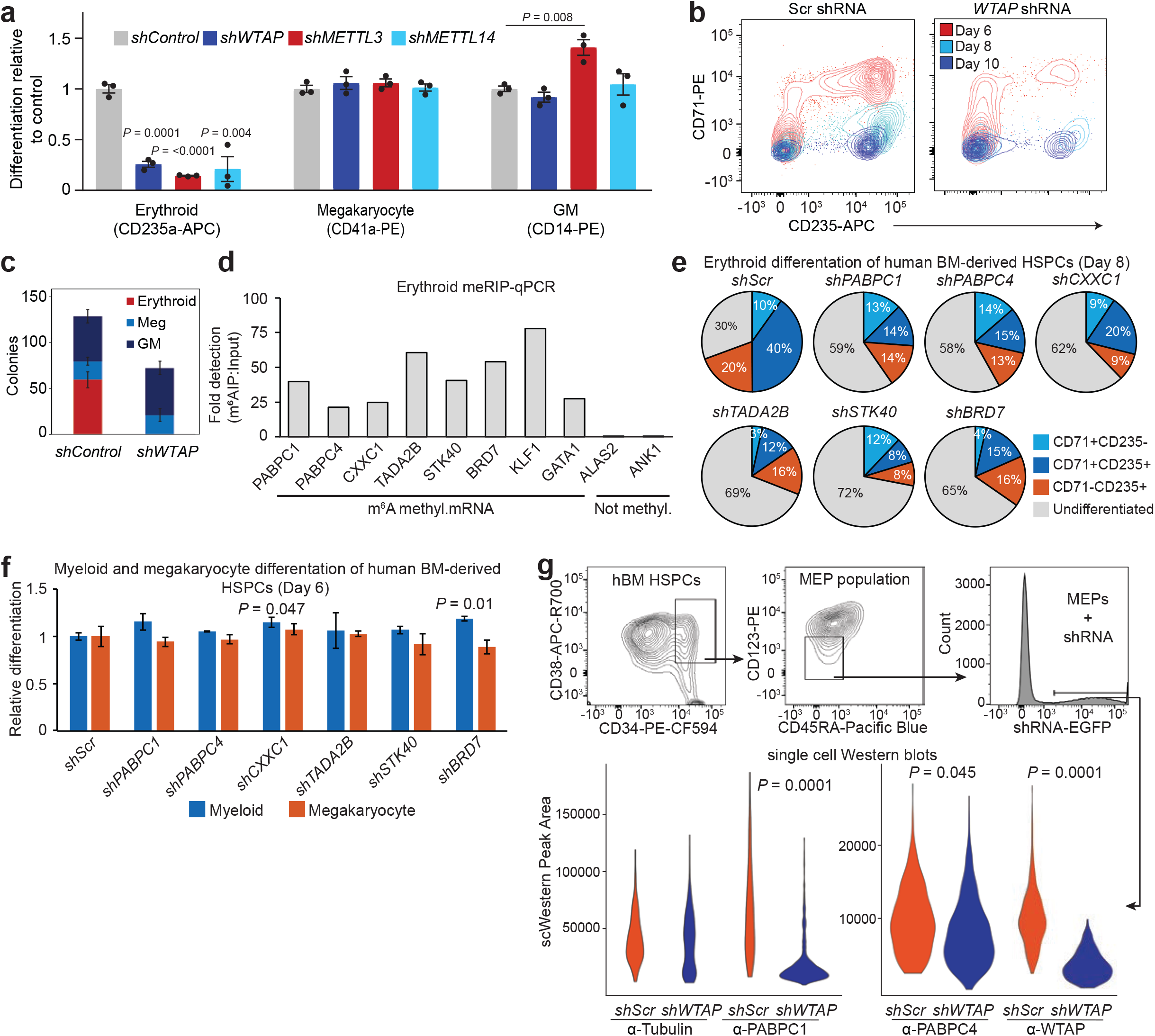
Knockdown of the m^6^A MTase complex blocks erythropoiesis in human adult, bone marrow-derived HSPCs. **a,** Flow cytometry of *WTAP* KD, *METTL3* KD and *METTL14* KD hBM HSPCs differentiated in liquid culture reveals a block to erythropoiesis with no impact on megakaryopoiesis and an enhancement of myelopoiesis with *METTL3* KD, (n=3, mean ± SEM, t-test 2-sided). **b,** Flow cytometry of erythroid progenitor populations reveals an early block to erythropoiesis following *WTAP* KD. At day 6, *WTAP*-KD CD235+/CD71+ cell numbers were only 8.4% of scr control and by day 10, *WTAP*-KD CD235+/CD71-cell numbers decreased further to 5.5% of control cells. **c,** Colony formation assays from *WTAP* KD hBM HSPCs confirm the block to erythropoiesis (n=6, mean ± s.d., erythroid t-test 2-sided p<0.0000001). **d,** meRIP-qPCR validation of m^6^A methylation for key erythroid regulators and genes translationally decreased following *WTAP* KO in a bulk population of adult BM erythroid cells (ERY1, ERY2 and ERY3) pooled from 6 donors. (n=1) **e,** Flow cytometry of lv-shRNA-KD of the translationally altered genes identified in Fig. 4c in hBM HSPCs differentiated in erythroid promoting liquid culture reveals a delay or reduction in erythropoiesis at day 8. (n=2, mean) **f,** Flow cytometry for differentiation potential of hBM CD34+ HSPCs transduced with lv-shRNA of the translationally altered genes identified in Fig. 4c and cultured in megakaryocyte or myeloid liquid culture differentiation conditions shows no effect on these lineages, except for enhanced myelopoiesis following *CXXC1*-KD and *BRD7*KD, (n=2, mean ± range, t-test 2-sided). **g,** Single cell Western blot analysis of WTAP-KD MEPs recapitulates the reduction in PABPC1 and PABPC4 protein observed in WTAP-KO HEL cells (data from a minimum of 250 cells were used per sample, Kolmogorov-Smirnov)

### Validation of m^6^A mRNA regulation targets in human adult, bone marrow-derived HSPCs

We next wished to determine whether key targets of m^6^A mRNA translational regulation in HEL cells, including *BRD7*, *CXXC1*, *PABPC1*, *PABPC4*, *STK40*, and *TADA2B*, show similar regulation in human erythroid progenitor populations and also have causal roles in promoting CD235a/GYPA expression and erythroid lineage formation. To this end, we first examined their m^6^A-mRNA status using RNA isolated from “bulk” erythroid progenitor populations, consisting of CD34-CD71+CD235a-, CD34-CD71+CD235a+ and CD34-CD71+CD235a+ cells and performed MeRIP-qPCR. We included genes which are not m^6^A marked in HEL cells, *ALAS2* and *ANK1*, as well as examples of key erythroid transcription factors which are m^6^A marked in HEL cells, *GATA1* and *KLF1* (Fig. 6d). By this approach, we detected significant m^6^A signal from *BRD7*, *CXXC1*, *PABPC1*, *PABPC4*, *STK40*, and *TADA2B* (21.2-fold to 60.9-fold enrichment versus total RNA) and *GATA1* and *KLF1* (27.7-fold and 78.4-fold enrichment, respectively), with minimal background detection of the unmethylated targets (0.01-fold to 0.10-fold detection versus total RNA) (Fig. 6d).

Next, we utilized lv-shRNAs to KD *BRD7*, *CXXC1*, *PABPC1*, *PABPC4*, *STK40*, and *TADA2B* in CD34+ adult BM cells and then assessed their impact on hematopoietic differentiation in liquid culture. KD of each of the genes, but not controls, strongly delayed the emergence of erythroid progenitors (~3 days), without negatively affecting megakaryocyte or monocyte lineage formation (Fig. 6e,f). Furthermore, none of the KDs produced significant differences in megakaryocytic differentiation, and only KD of CXXC1 and BRD7 produced a small but significant increase in myeloid differentiation (Fig. 6f). These results demonstrate that key m^6^A-MTase mRNA targets are also found to be m^6^A-regulated in primary erythroid progenitors and that their partial inhibition blocks or delays human erythroid lineage specification.

### *WTAP* is required for *PABPC1* and *PABPC4* protein expression in human megakaryocyte-erythroid progenitors

We next tested whether m^6^A-mRNA regulation may affect translation regulation of *BRD7*, *CXXC1*, *PABPC1*, *PABPC4*, *STK40*, and *TADA2B* in human HSPCs. However, because m^6^A-MTase inhibition blocks the appearance of erythroid progenitors in human HSPCs and due to the paucity of cells in progenitor populations from human donors (e.g., traditional Western-blotting is not possible), we examined the effects of *WTAP* KD on expression in flow sorted MEP populations using single-cell Western blotting, where only hundreds of single cells are in theory needed to analyze protein expression (Supplementary Fig. 6d). Unfortunately, only PABPC1 and PABPC4 proteins were detectable at measurable levels using this assay (Fig. 6g and Supplementary Fig. 6e); so, we proceeded testing only them. Both *PABPC1* and *PABPC4* encode cytoplasmic poly A binding proteins shown to promote the expression of β-globin in erythroid-differentiated CD34+ cells^68^ and promote the erythroid maturation of mouse erythroid leukemia cells^53^, respectively. By this analysis, targeted lv-based KD of *WTAP* in MEP-enriched populations caused significant reduction in both PABPC1 and PABPC4 protein, but not control beta-tubulin. We conclude that WTAP activity promotes their protein expression in human HSPCs and likely does so through m^6^A-mRNA regulation (Fig. 6g).

## Discussion

Here, we report the discovery of a new form of post-transcriptional regulation of human erythroid gene expression and lineage specification, m^6^A-mRNA marking. From whole-genome CRISPR-Cas9 screening, we found that the three core components of the m^6^A MTase complex (*METTL14*, *METTL3*, and *WTAP*) are required for maintaining CD235a/GYPA expression in HEL cells and erythroid lineage specification in human HSPCs. In depth mapping of the m^6^A mRNA methylome in HEL and primary hematopoietic cell populations revealed that many key hematopoietic and erythroid factors are subject to m^6^A-mRNA marking, including, for example, *GATA1*, *GATA2*, *KLF1*, *RUNX1*, and *SPI1* (Fig. 2). Consistent with m^6^A-MTase activity playing key roles in promoting erythroid gene expression, we observe that KO of *METTL14*, *METTL3*, and *WTAP* caused loss of erythroid transcription programs (Fig. 3).

Mechanistically, we found that m^6^A mRNA regulation promotes the translation of hundreds of genes, many of which have causal roles in erythropoiesis and erythroid-associated diseases or appeared in two prominent gene networks: histone H3K4 MTases and interacting proteins, and ribosome/RNA binding proteins. Comprehensive validation of m^6^A mRNA targets translationally regulated in HEL cells revealed that *BRD7*, *CXXC1*, *NIP7*, *PABPC1*, *PABPC4*, *RPS19*, *RPL24*, *STK40*, and *TADA2B* are required for promoting CD235a/GYPA expression (Fig. 4). Further, each contained a 197-438 bp mRNA m^6^A regulator element which could be transferred to luciferase reporter gene and impart translation enhancement (Fig. 4).

Remarkably, follow up studies on m^6^A-mRNA-dependent regulation of SETD1A/B H3K4 MTase activity revealed that m^6^A-MTase activity was required for maintaining H3K4me3 marks for many erythroid-related genes, including those involved in the KLF1 transcriptional program, in particular, the heme and hemoglobin synthesis pathway (Fig. 5). Taken together, the results demonstrate that m^6^A-mRNA plays key roles in promoting human erythroid gene expression and lineage specification (Fig. 5f).

Previous work has established three prominent types of post-transcriptional mRNA regulation impacting erythropoiesis or expression of key erythroid genes. These include: microRNA-based targeting of critical pro- or anti-erythroid transcription factors (rev. in^69^); erythroid-specific messenger ribonucleoprotein-dependent regulation of *βGlobin/HBB* mRNA stability^68^; and the iron regulatory protein/iron-responsive element regulatory system, which post-transcriptionally controls the translation of several key heme biosynthesis genes^70,71^. Our results provide insight into another critically, yet unexpected, layer of post-transcriptional mRNA regulation in erythroid cells with several points of intersection with these other forms of regulation. For example, one of the key m^6^A-mRNA targets in HEL cells and in HSPCs, PABPC1, is recruited by messenger ribonucleoprotein onto *β-Globin/HBB* mRNA to prevent its deadenylation and decay^68^. We also find that m^6^A-MTase activity promotes both steady-steady state mRNA levels and H3Kme4 marks of many heme biosynthesis genes, including ones with IREs (e.g., TRFC and ALAS2) (Fig. 5d).

Our work also sheds light on how epitranscriptomic m^6^A regulation can impact epigenetic patterning of chromatin. We unexpectedly found that m^6^A-MTase activity controls translation of a network of histone H3K4 MTases, including SETD1A, SETD1B, and KMT2D MTases and their associated proteins (Fig. 4). Consistent with this notion, we found that H3K4me3 marks are dramatically reduced after *METTL14*, *METTL3*, and *WTAP* KO at sites of KLF1 binding (Fig. 5). SETD1A and KMT2D have previously been shown to have key roles in regulating erythroid gene transcription, and, at least for SETD1A, key roles in erythroid lineage specification^55,56^. Our results in HEL cells suggest that m^6^A marks may play a key role in maintaining optimal H3K4me3 marking and, as a result, proper expression of lineage genes.

Phenotypically, in adult human HSPCs, our studies find that m^6^A MTase function is critical just after the MEP stage of erythroid lineage specification. Intriguingly, while this erythropoiesis failure occurs at an earlier stage than observed in Diamond Blackfan Anemia (DBA) and myelodysplastic syndrome (MDS)^5^, a number of the key drivers for these diseases are m^6^A targets and require m6A-MTAase activity for their expression. Loss of these gene activities may explain the maturation failure of the small numbers of detected early erythroid progenitors (Fig. 6). We detected m^6^A mRNA methylation in 70 out of 104 known/putative MDS genes, including 8 of the top 10 most frequently mutated genes (*TET2*, *SF3B1*, *ASXL1*, *RUNX1*, *DNMT3A*, *ZRSR2*, and *STAG2*)^72^. Further, *RPS19*, the most commonly mutated gene in DBA (~25%) is decreased translationally following *WTAP* KO, as well as less frequently mutated *RPS10* (~2.6%)^73^. This could suggest that m^6^A mRNA regulation could emerge as an erythroid-associated disease modifier.

Additionally, we validated roles for multiple m^6^A-mRNA regulatory targets in CD235a/GYPA expression and for erythroid lineage specification in hHPSCs, including: *BRD7*, *CXXC1*, *NIP7*, *PABPC1*, *PABPC4*, *RPL24, RPS19*, *STK40*, and *TADA2B*. Of these, NIP7 has been previously implicated in 18S rRNA maturation and the MDS-associated Shwachman-Bodian-Diamond syndrome^74^. *PABPC1* and *PABPC4* encode polyA RNA binding proteins. *PABPC1* has been shown to bind to and stablize the *βGlobin/HBB* mRNA by inhibiting its deadenylation in erythroid precursors^68^, whereas *PABPC4* has been implicated in binding to and stabilizing *GYPA* mRNA, as well as other erythroid targets, including *α-Globin/HBA1/HBA2*, *β-Globin/HBB*, *BTG2*, and *SLC4A1*^53^. *RPS19* encodes a ribosomal protein that when heterozygous for a loss of function mutation causes DBA^73^. *STK40* codes for a serine threonine kinase required for definitive erythropoiesis just after the MEP state in the fetal mouse liver, similar to our m^6^A-MTase inhibition phenotype in hHSPCs^54^. The other genes have not been previously implicated in erythropoiesis, including: *BRD7*, a member of the bromodomain-containing protein family implicated in tumor suppression of p53 and PI3K pathways^75,76^; *CXXC1*, which encodes a key regulator of H3K4 histone methylation, cytosine methylation, cellular differentiation, and vertebrate development^77^; and *TADA2B*, which encodes a transcriptional adaptor for transcriptional activation factors^78^.

Taken together, our results reveal a new physiological role for m^6^A-dependent regulation of mRNA translation in maintaining erythroid gene expression programs and promoting erythroid lineage specification and provide a key resource for the study of epitranscriptomics during human hematopoiesis and erythropoiesis.

## Methods

### Cloning

#### sgRNAs

*SgRNAs* were cloned into lentiCRISPRv2 puro vector (Addgene) and lentiCRISPRv2-mCherry (a variant created by excising puro with NheI and MluI, followed by addition of P2A-mCherry by Gibson Assembly). Individual gene pools of 4 sgRNAs were cloned by first PCR amplifying the individual sgRNAs from oligo template using Phusion High-Fidelity DNA Polymerase (NEB), followed by cleanup with the PureLink PCR Purification Kit (Invitrogen). The following primers were used for sgRNA amplification: Array_F-TAACTTGAAAGTATTTCGATTTCTTGGCTTTATATATCTTGTGGAAAGGACGAAACA CCG and Array_R-ACTTTTTCAAGTTGATAACGGACTAGCCTTATTTTAACTTGCTATTTCTAGCTCTAAAAC. The oligo template is a 60-mer oligo: GTGGAAAGGACGAAACACCg-sgRNA sequence – GTTTTAGAGCTAGAAATAGC. The individual sgRNA PCR products were quantified on a Nanodrop 1000 (ThermoFisher) and pooled in equal amounts. The PCR product pools were then cloned into the Esp3I site of lentiCRISPRv2 by Gibson Assembly. Four colonies from each pool were sequenced to validate the pool identity, while the remainder of the plate was scraped and DNA prepared using the NucleoBond Xtra kit (Macherey-Nagel).

#### shRNAs

A modified pLL3.7 vector (pLL3.7-EF1a-mini) was generated replacing the CMV promoter with EF1a and inserting an Esp3I cloning cassette. The CMV promoter was removed by NsiI and AgeI digest and the EF1a promoter inserted by Gibson Assembly. A new shRNA cloning site was added by digesting with XhoI and NsiI and inserting a 100-bp cassette containing Esp3I sites on both ends by Gibson Assembly. All shRNAs were cloned into the pLL3.7-EF1a-mini vector, primarily following the Genetic Perturbation Platform shRNA/sgRNA Cloning Protocol, (portals.broadinstitute.org/gpp/public/resources/protocols) with one modification Due to the altered cloning site, the format for the annealed oligos changed to the follow: 5’ GTTT---21 bp Sense --- CTCGAG --- 21 bp Antisense --- TTTTTG 3’ and 5’ GCCGCAAAAA --- 21 bp Sense --- CTCGAG --- 21 bp Antisense --- 3’. The vector is linearized with Esp3I and gel purified using the Monarch Gel Extraction Kit (NEB) or Zymoclean Gel DNA Recovery Kit (Zymo Research). All sgRNA and shRNA sequences can be found in Table S8.

#### Luciferase reporters

Regions of potential m^6^A regulation were cloned into the pPIG vector (this paper). The m6A regions of interest were ordered as either WT or mutant, with all central adenines within the m6A methylation motif mutated, gBlocks (IDT). Luciferase was PCR amplified out of Luciferase-pcDNA3. Luciferase and the reporter region were cloned between the NotI and MluI sites of pPIG by Gibson Assembly. pPIG is heavily modified version of pGIPZ with Hygro resistance removed and the region between the XbaI and MluI sites replaced with the following cassette, XbaI – hPGK promoter – NotI – IRES – AgeI – EGFP – MluI.

### Cell Culture and lentiviral transduction

HEL cells were grown in RPMI-1640 (ThermoFisher) supplemented with 10% fetal bovine serum (FBS) and maintained a concentration between 250,000 and 1 million cells/ml. 293T cells were grown in Dulbecco’s modified Eagle’s medium (ThermoFisher) supplemented with 10% fetal bovine serum. For lentiviral transduction, HEL cells were plated at 250,000 cells/ml in RPMI-1640 + 10% FBS, 8 μg/ml protamine sulfate (Fisher Scientific ICN19472905) and virus is added at the MOIs described below. After 48 hours, the cells are washed with 1× PBS and fresh RPMI + 10% FBS added.

### Virus production

The lentiCRISPRv2 vector was used for all CRISPR-Cas9 experiments and the pLL3.7EF1a vector was used for all RNAi experiments. For virus production, 12 μg of either lentiCRISPRv2 or pLL3.7-EF1a, 8 μg of psPAX and 3 μg of pMD2.G were transfected with PEI into 293T cells. Approximately 24 hours post transfection the media was replaced with fresh DMEM containing 10% FBS. Viral supernatants were harvested and filtered 24 hours later and immediately concentrated with a 20-24 hours spin at 6000 ×g. Approximately 100× concentrated virus was stored at −80C.

### CRISPR-Cas9 Screening

The Human GeCKOv2 whole genome library^79^ was used in lentiviral pooled format to transduce HEL cells. For each screen replicate, cells were transduced at approximately 1000-fold representation of the library (at 30% infection efficiency). 2 days after transduction, 2 ug/ml puromycin was added for 3 days. A portion of cells representing 500-fold coverage of the library were harvested as the day 0 timepoint. The rest of the cells were then passaged to maintain 500-fold representation and cultured for an additional 7 days. Genomic DNA was extracted, and a previously described two-step PCR procedure was employed to amplify sgRNA sequences and then to incorporate deep sequencing primer sites onto sgRNA amplicons. Purified PCR products were sequenced using HiSeq 2000 (Illumina). Bowtie^80^ was to align the sequenced reads to the guides. The R/Bioconductor package edgeR^81^ was used to assess changes across various groups. Guides having a fold change more than 1 and an adjusted FDR <0.05 were considered statistically significant.

### CRISPR and Ribosome profiling retests

For individual gene retests of the initial whole genome CRISPR-Cas9 screen, 4 sgRNA pools were cloned, as described above. The sgRNAs were selected by combining the sgRNAs which scored in the primary screen with additional sgRNAs from the human CRISPR Brunello lentiviral pooled library. 2×10^5^ HEL cells were transduced at MOI 10 and CD235a expression was measured by flow cytometry after 10 days.

Ribosome profiling individual retest pools for non-essential genes contained 3 sgRNAs, but were otherwise cloned and tested as described above for the CRISPRCas9 screen. For essential genes shRNAs were selected from the Genetic Perturbation Platform database (portals.broadinstitute.org/gpp/public/) based on those with the highest adjusted score. Three shRNA were used per gene and tested individually. 2×10^5^ HEL cells were transduced at MOI 10 and CD235a expression was measured by flow cytometry after 4-7 days.

### Adult bone marrow CD34+ cell isolation, culture

Bone marrow aspirates were collected under FHCRC IRB protocol 0999.209. The cells isolated from aspirates were considered non-human subjects as no identifiable information was associated with the leftover specimen. The collected aspirates were washed twice in 1× PBS, Ficoll fractionated and the mononuclear cell fraction collected. Enrichment for CD34+ cells was done by magnetic bead isolation using the CD34 MicroBead Kit (Miltenyi Biotec). CD34+ HSPCs were grown in StemSpan SFEM II (Stemcell Technologies) supplemented with the following growth factor cocktails: expansion (SCF 100 ng/ml, FLT-3 ligand 10ng/ml, TPO 100 ng/ml, IL-6 20 ng/ml), erythroid (SCF 20 ng/ml, IL-3 10 ng/ml and EPO 4 U/ml), myeloid (SCF 100 ng/ml, FLT3 ligand 10 ng/ml, IL-3 20 ng/ml, IL-6 20 ng/ml, GM-CSF 20 ng/ml, M-CSF 20 ng/ml and G-CSF 20 ng/ml), megakaryocyte (SCF 10 ng/ml, IL-6 20 ng/ml, IL-9 12.5 ng/ml and TPO 100 ng/ml). All cytokines were purchased from Shenandoah Biotechnology.

### Adult bone marrow CD34+ cell transduction

CD34+ cells were thawed and cultured in StemSpan SFEM II + expansion growth factor cocktail for 16h prior to transduction. Plates were then coated with 10 μg/cm2 RetroNectin (Takara) for 2 hours at RT, following the manufactures instructions, followed by preloading with virus (final MOI 50) for 15 mins at RT. hBM CD34+ cells were then cultured on the plates for 48 hours at 500,000 cells/ml in StemSpan SFEM II + expansion growth factor cocktail. The cells were then washed twice with 1× PBS and re-plated in StemSpan SFEM II + expansion growth factor cocktail at 250,000/ml.

### Flow cytometry

HEL cells were stained with APC-CD235a (BD Pharmingen 551336). To monitor CD34+ progenitor cell differentiation status the cells were stained with PE-CF594-CD34 (BD Pharmingen 562383), APC-R700-CD38 (BD Pharmingen 564979), PE-CD123 (BD Pharmingen 554529), Pacific Blue CD45RA (Invitrogen MHCD45RA28), APC-H7-CD41 (BD Pharmingen 561422) and BV786-CD71 (BD Pharmingen 563768). To monitor lineage status cells were stained with APC-CD235a (BD Pharmingen 551336) and PECD71(BD Pharmingen 555537) for erythropoiesis, PE-CD14 (BD Pharmingen 555398) for myeloid cells, and PE-CD41 (BD Pharmingen 557297) and APC-CD61 (BD Pharmingen 564174) for megakaryopoiesis. Cells were analyzed on an BD LSRII instrument.

### CFU assays

10^4^ CD34+ shWTAP transduced cells were plated for lineage specific colony assays (triplicate) in methylcellulose (MethoCult H4230, Stem Cell Technologies) supplemented with the lineage specific growth factor cocktails outlined above. Colonies were scored after 14 days in culture.

### meRIP-seq

meRIP-seq was performed largely as previously described ^36^. Total RNA from HEL cells was isolated by Trizol (ThermoFisher) and the Direct-zol RNA kit (Zymo Research). Poly(A) RNA was then isolated with the NucleoTrap mRNA Mini kit (Macherey-Nagel) yielding approximately 5 μg of poly-A RNA per replicate. The poly-A RNA was then fragmented using the NEBNext Magnesium RNA Fragmentation Module (NEB) for 7 mins at 94C yielding RNA fragments of approximately 125 bp. Fragmented RNA was incubated with 5 μg m6A antibody (Millipore Sigma) for 2 hours, followed by immunoprecipitation and phenol:chloroform cleanup and ethanol precipitation. Sequencing libraries were generated using the KAPA Biosystems Stranded RNA-Seq Kit (Roche), were quantified using a Qubit Fluorometer (Thermo Fisher), and the size distribution was checked using TapeStation 4200 (Agilent Technologies). 50 bp paired-end reads were sequenced on an Illumina HiSeq 2000.

### Primary cell meRIP-seq

Total RNA from HEL cells of flow sorted primary hematopoietic cell populations was isolated by Trizol (ThermoFisher) and the Direct-zol RNA kit (Zymo Research). The concentration of total RNA was measured by Qubit RNA HS Assay Kit (ThermoFisher). A total of 3 μg of total RNA was then fragmented using the NEBNext Magnesium RNA Fragmentation Module (NEB) for 6 mins at 94C yielding RNA fragments of approximately 150 bp. The fragmented RNA was precipitated overnight at −80C and resuspended in 10ul H_2_O per 1ug of total RNA. Per sample 30 μl of protein-A magnetic beads (NEB) and 30 μl of protein-G magnetic beads (NEB) were washed twice with IP buffer (150 mM NaCl, 10 mM Tris-HCl, pH 7.5, 0.1% IGEPAL CA-630 in nuclease free H_2_O), resuspended in 250 μl of IP buffer, and tumbled with 5 μg anti-m6A antibody at 4°C for at least 2 hrs. Following 2 washes in IP buffer, the antibody-bead mixture was resuspended in 500 μl of IP reaction mixture containing the fragmented total RNA (minus 50 ng of input control RNA), 100 μl of 5× IP buffer and 5 μl of RNasin Plus RNase Inhibitor (Promega), and incubated for 2 hrs at 4°C. The RNA reaction mixture was then washed twice in 1 mll of IP buffer, twice in 1 ml of low-salt IP buffer (50 mM NaCl, 10 mM Tris-HCl, pH 7.5, 0.1% IGEPAL CA-630 in nuclease free H2O), and twice in 1 ml of high-salt IP buffer (500 mM NaCl, 10 mM Tris-HCl, pH 7.5, 0.1% IGEPAL CA630 in nuclease free H_2_O) for 10 min each at 4°C. After washing, the RNA was eluted from the beads in 200 μl of RLT buffer supplied in RNeasy Mini Kit (QIAGEN) for 2 min at room temperature. The magnetic separation rack was applied to pull beads to the side of the tube and the supernatant was collected to a new tube. 400 μl of 100% ethanol was added to the supernatant and RNA isolated following the manufactures instructions and eluted in 14 μl. 2 μl of eluted RNA was reverse transcribed with SuperScript IV Reverse Transcriptase (Thermo Fisher) and IP efficiency was assessed by KLF1/GAPDH qPCR in a HEL control sample. 3.5 μl of eluted RNA and 50 ng of input RNA were used for library construction with the SMARTer^®^ Stranded Total RNA-Seq Kit v2 - Pico Input Mammalian (Takara) according to the manufacturer’s protocol. Libraries for IP RNA were PCR amplified for 16 cycles whereas 12 cycles were used for input RNA. The purified libraries were quantified using a Qubit Fluorometer (Thermo Fisher), and the size distribution was checked using TapeStation 4200 (Agilent Technologies) and 50 bp paired-end reads were sequenced on an Illumina HiSeq 2500.

### meRIP-qPCR

m^6^A RNA was isolated as described for primary cell meRIP-seq without the fragmentation step. Random hexamers and SuperScript IV Reverse Transcriptase (Thermo Fisher) were used to generate cDNA and qPCR run with Power SYBR Green Master Mix (Thermo Fisher) on a QuantStudio 7 Flex (Thermo Fisher).

### meRIP-seq analysis

50bp paired end sequenced reads were mapped to hg19 using TopHat v2 ^82^ and the resulting bam files were processed using Picard tools (http://broadinstitute.github.io/picard) to mark duplicate reads. Custom R scripts were written for identification of peaks based on ^37^ and for visualization of peaks. The Human transcriptome (from hg19) was broken into 25nt wide discrete non-overlapping reads. Using bedtools ^83^ the 50bp reads were mapped to 25nt windows and windows with fewer than 5 aligned reads were dropped. We used one sided Fisher’s exact test to compare the number of reads that mapped to a given window for the MeRIP sample and the non-IP sample to the total number of reads in each. The Benjamini-Hochberg procedure was used to adjust the p-values from the Fisher’s exact test to reduce our false discovery rate to 5%. To find a final p-values for each window, Fisher’s Method was used to combine p-values across replicates. To find distinct m6A peaks, we combined consecutive significant 25nt windows across the transcriptome. Consecutive significant windows less than 100bp were discarded. Gene region annotations for the peaks were found from UCSC RefSeq Table for Hg19 using R/Bioconductor package TxDb.Hsapiens.UCSC.hg19.knownGene (https://bioconductor.org/packages/release/data/annotation/html/TxDb.Hsapiens.UCSC.hg19.knownGene.html). Motif calling was done using HOMER (http://homer.ucsd.edu/homer/motif/) and the default settings.

### RNAseq and splicing analysis

HEL *sgGYPA*-KO, *sgGATA1*-KO, *sgLMO2*-KO, and *sgWTAP*-KO cells were lysed with Trizol (ThermoFisher) Total RNA was isolated with the Direct-zol RNA kit (Zymo Research) and quality validated on the Agilent 2200 TapeStation. Illumina sequencing libraries were generated with the KAPA Biosystems Stranded RNA-Seq Kit (Roche) and sequenced using HiSeq 2000 (Illumina). RNA-seq reads were aligned to the UCSC hg19 assembly using Tophat2 ^82^ and counted for gene associations against the UCSC genes database with HTSeq ^84^. The normalized count data was used for subsequent Principal component analysis and Multidimensional scaling (MDS) in R. Differential Expression analysis was performed using R/Bioconductor package DESeq2 ^85^.

The Miso pipeline ^41^ was used to analyze the RNA-seq data for alternatively spliced transcripts. First, the expression levels (psi values) were computed for each of the paired end RNA-seq samples individually using ‘miso --run’, followed by calculating the Psi values for each sample using ‘summarise-miso’. Lastly, all pairwise comparisons between the *sgNTC* and *sgWTAP* samples were run using ‘compare_miso’ and the events were fitered using criteria ‘--num-inc 1 --num-exc 1 --num-sum-inc-exc 10 --deltapsi 0.10 --bayes-factor 10’. Only those alternative splicing events that were present in all pairwise comparisons were included in our final results.

### m^6^A colorimetric quantification

The manufactures instructions for the m^6^A RNA Methylation Quantification Kit (Abcam) were followed. HEL cells transduced with *sgWTAP* and *sgGATA1* were flow sorted for CD235a^-/low^ expression. RNA was isolated as described above for RNAseq and 200 ng of total RNA used per assay. Samples were assayed in triplicate.

### m^6^A luciferase reporters

HEL cells were transduced in triplicate with the lv-m^6^A-luciferase reporters at an MOI 10 as described above and cultured for 4 days. The cells were then collected, and half used for detection of luciferase with the Dual-Luciferase Reporter System (Promega) following the manufactures instructions and luminescence measured on a GloMax Multi+ (Promega). Genomic DNA was extracted from the other half using the E.Z.N.A. MicroElute Genomic DNA Kit (Omega Bio-tek). The transduction efficiency of each sample was then quantified by WPRE qPCR as previously described ^86^ using Power SYBR Green Master Mix (Thermo Fisher) on a QuantStudio 7 Flex (Thermo Fisher). The luciferase signal was then normalized to the transduction efficiency.

### Ribosome profiling and analysis

The profiling methodology was based largely on protocols described by Ingolia and Hsieh and colleagues ^42,43^. Wild-type and *WTAP* knockdown HEL cells were utilized to generate both ribosome protected RNA libraries as well as alkaline digested total RNA libraries using the ARTseq Ribosome Profiling Kit (Illumina). Ribo-Zero (Illumina) was used to subtract rRNA levels from each replicate. Once samples were generated, they were sequenced using an Illumina HiSeq 2500. The raw sequence data was uncompressed followed by clipping the 3’ adaptor sequence (AGATCGGAAGAGCACACGTCT). Next, trimmed sequence reads were aligned to an rRNA reference using Bowtie ^80^. rRNA alignments were removed to reduce rRNA contamination. The unaligned reads were collected and TopHat2 ^82^ was utilized to align non-rRNA sequencing reads to hg19. Reads for each gene were counted using HTSeq (UCSC reference transcriptome) ^84^. Reads were only counted starting from 20 nucleotides after the start codon and up to 20 nucleotides before the stop codon. R/Bioconductor package, DESeq2 ^85^ was used to identify differentially expressed genes at the translational level using both ribosome bound and total RNA samples. A statistical cutoff of FDR < 0.05 and log2 fold change > 1 was applied to find translationally and transcriptionally regulated genes. RiboseqR ^87^ was used to calculate triplate periodicity in all samples. Translation efficiency is a measure of ribosome bound mRNA over total mRNA and is a snapshot of the translation rate of an mRNA of interest. R/Bioconductor package, edgeR ^88^ was used to find differentially expressed gene at the transcriptional level using only rRNA subtracted total RNA specimens.

### CUT&RUN and analysis

CUT&RUN was performed as previously described ^89^. 200,000 HEL cells were harvested and bound to Concanavalin A–coated beads at RT for 10 mins. The bound cells were permeabilized with 0.025% Digitonin with a 1:50 dilution of the H3K4me3 primary antibody (Cell Signaling Technologies), 1:100 dilution of H3K27me3 primary antibody (Cell Signaling Technologies) as a positive control and, 1:50 dilution of Guinea Pig anti-Rabbit IgG (Antibodies-Online) as a negative control, followed by incubation with rotation overnight at 4°C. The antibody bound cells were incubated with the pA-MNase at a 1:10 dilution for 1 hour at 4°C and the MNase digestion was then run for 30 mins at 0°C. Released DNA fragments were extracted using the NucleoSpin Gel and PCR Clean-up kit (Macherey-Nagel) as described by Skene et al. 2018 and sequencing libraries generated using the KAPA HyperPlus kit (Roche) following the manufactures instructions.

Sequencing was performed using an Illumina HiSeq 2500 in Rapid Run mode and employed a paired-end, 50 bp read length (PE50) sequencing strategy. Raw fastq files were aligned with Bowtie ^80^ to the human genome (hg19) and spike in to the Saccharomyces cerevisiae genome(sacCer3), followed by spike in calibration. Picard (https://broadinstitute.github.io/picard/) was used to remove duplicate reads and aligned fragments were extracted from the bam file to a bed file. The bed file was used to call peaks following the protocol described by Skene et al. ^89^. EaSeq was used for data visualization ^90^.

### Western blot and puromycin incorporation

Immunoblots were performed following standard protocols (www.cshprotocols.org). HEL cells were lysed in a modified RIPA buffer (150mM NaCl, 50mM Tris, pH 7.5, 2mM MgCl2, 0.1% SDS, 2mM DDT, 0.4%Triton X-100, 1X complete protease inhibitor cocktail (complete Mini EDTA-free, Roche) on ice for 15 mins. Cell lysates were quantified using the Pierce BCA Protein Assay Kit (Thermo Fisher). The Trans-Blot Turbo Trans-Blot Turbo transfer system was used according to the manufacturer’s instructions. The following commercial antibodies were used: CXXC1 (Cell Signal, 1:250), PABPC4 (Novus Biologicals, 1:500), PABPC1 (Thermo Fisher, 1:500), WTAP (Abcam, 1:500), BRD7 (Thermo Fisher, 1:250), STK40 (Thermo Fisher, 1:250), TADA2B (Abnova, 1:250), Beta-actin (Cell Signaling, 1:1,000), and Beta-tubulin (Abcam, 1:1,000). An Odyssey infrared imaging system (LI-COR) was used to visualize blots following the manufacturer’s instructions. The Image Studio software was used to semi-quantify the blots.

The puromycin incorporation assay was performed as previously reported^91^. HEL cells were transduced with *sgNTC* or sg*WTAP* and cultured for 10 day. On day ten, the cells were treated with 10 ug/ml of puromycin for 10 mins at 37C. Immunoblotting for puromycin incorporation (Millipore Sigma, 1:1000) was done as described above.

### Single-cell Western blotting^92^

Adult bone marrow CD34+ HSPCs were transduced with *shScr* or *shWTAP* lentivirus and cultured in the expansion growth factor cocktail as described above. Five days post transduction the cells were flow sorted for the transduced (GFP+) MEP progenitor population as shown in Fig. 6g. The sorted cells were then immediately loaded onto the Milo small scWest Chip (ProteinSimple) following the manufactures instructions. The following run conditions were used: Lysis – 5 seconds; Electrophoresis – 70 seconds; UV Capture 4 minutes. The chips were probed with the following antibodies: PABPC4 (Novus Biologicals, 1:20), PABPC1 (ThermoFisher, 1:20), WTAP (Abcam, 1:20), Betatubulin (Abcam, 1:20) and Alexa Fluor 555 donkey anti-rabbit (Invitrogen, 1:40) was used for the secondary antibody. Chips were scanned on a GenePix 4000B (Molecular Devices) and analyzed with the scout software package (ProteinSimple).

### Gene set enrichment analysis

Genes sets arising from our genomic data sets were analyzed using GSEA^93^, the ToppGene tool suite^94^, or GeneMANIA network viewer plugin for Cytoscape^95,96^.

### Reporting summary

Further information on experimental design is available in the Nature Research Reporting Summary linked to this article.

### Code availability

m^6^A peak calling was performed using custom R scripts that will be provided upon request. All other analyses were performed using publicly available software as indicated.

### Data availability

sgRNA-seq, RNA-seq, Ribo-seq, CUT&RUN and MeRIP-seq raw reads and processed data sets can be accessed at NCBI Gene Expression Omnibus under accession number GEO: GSE106124.

## Acknowledgements

We thank members of the Paddison, Hsieh, and Torok-Storb labs for helpful suggestions, the Henikoff lab for Cut & Run protocols and advice, Pam Lindberg and Lori Blake for administrative support, the Fred Hutch NIDDK-CCEH Cell Processing Core for providing aspirated bone marrow, and Andrew Marty at the Fred Hutch Genomics Shared Resource for technical assistance with and advice for deep-sequencing runs. This work was supported by the following grants: DOD PCRP Postdoctoral Training Award (PC150946)(YL), an AACR-Bristol-Myers Squibb Fellowship (Y.L.), a V Foundation Scholar Award (ACH), the AACR NextGen Grant for Transformative Cancer Research (ACH), NIH 1K08CA175154-01 (ACH), the Burroughs Wellcome Fund Career Award for Medical Scientists (ACH), NHLBI-U01 HL099993-01 (BTS & PJP), NHLBI-U01 Pilot project HL099997 (PJP), NIDDK-P30DK 56465-13 (BTS & PJP), NIDDK-U54DK106829 (BTS & PJP), and American Cancer Society Research Scholar Grant ACS-RSG-14-056-01 (PJP). This research was funded in part through the NIH/NCI Cancer Center Support Grant P30 CA015704.

## Contributions

D.K. and P.J.P. conceived of the initial idea and the screen. D.K., B.T-S., A.C.H., and P.J.P. designed follow up and mechanistic experiments. D.K. and A.L. performed experiments. Y.L., L.C., S.W., H.H.H. and P.C. provided technical assistance. S.A., R.B., and C.P. performed computation analyses and statistical tests. J.D. oversaw deep-sequencing runs and some of the primary data analysis. D.K., S.A., Y.L., A.C.H., and P.J.P. wrote and revised the manuscript.

